# Fibronectin deficiency in newborn mice leads to cyst formation in the kidney

**DOI:** 10.1101/2022.07.18.500556

**Authors:** Kristina Hermann, Silke Seibold, Kathrin Skoczynski, Bjoern Buchholz, Ernst R. Tamm, Leonie Herrnberger-Eimer

## Abstract

**Purpose:** The ubiquitously expressed glycoprotein fibronectin (FN) is a central component of the fibrillar extracellular matrix (ECM) that is found in multiple sites throughout the body including the peritubular interstitium of the kidney. To learn more about the specific role(s) of FN in the kidney we generated and investigated FN-deficient mice.

**Methods:** We generated CAGG-Cre-ER^™^/Fn^fl/fl^ mice which carry floxed Fn alleles and ubiquitously express Cre-recombinase after tamoxifen treatment. Newborn pups were treated with tamoxifen eye drops (2.5 mg/mL) to induce FN deficiency. Conditional deletion of Fn was confirmed by quantitative real-time PCR, Western blot analysis and immunohistochemistry. The expression patterns of Fn were analyzed by *in situ* hybridization. Kidneys were investigated by light microscopy and immunohistochemistry.

**Results:** The expression analyses and immunohistochemistry showed a significant reduction of FN at postnatal day (P) 4. Loss of FN corelated with the formation of renal cysts at the corticomedullary border, which expand with increasing age. *In situ* hybridization demonstrated that on P4 Fn expression extends mainly from the pelvis to the corticomedullary border, whereas in 5-6 weeks old mice it is located only in the cortex. Immunohistochemistry and light microscopy showed a loosening of the renal interstitium and additionally an appearance of ECM proteins in the cysts.

**Conclusion:** We conclude that FN deficiency leads to the development of renal cysts, which occurs a few days after tamoxifen treatment and results in extensive loss of renal parenchyma a few weeks after birth. The results indicate an important role of FN for maintenance of kidney structure and function.

## Introduction

Fibronectin (FN) is a 230–270 kDa glycoprotein and important constituent of the vertebrate extracellular matrix (ECM), which is ubiquitously expressed in human tissues.^1^ FN is encoded by a single gene and exists in multiple isoforms that are generated by alternative splicing. FN dimers are deposited from cells in a compact form into the ECM (cellular FN) or secreted by the liver into the plasma (plasma FN). Both forms then contribute to the generation of FN fibrils in the ECM. FN fibrillogenesis is a non-spontaneous cell-mediated process that involves the interaction with cell-surface integrins. There is evidence that the deposited FN matrix is essential for further formation and incorporation of other fibrillar matrix molecules such as collagens I and III in order to form the mature ECM.^2, 3^ FN is expressed early in the developing kidney^4^ and appears to be required to initiate and maintain branching morphogenesis during early embryonic formation of the urinary collecting duct system.^5, 6^

In the normal adult kidney FN is detected in the glomerular mesangium and the peritubular interstitium.^7^ Plasma FN contributes to mesangial FN, but not to interstitial FN which is largely of local origin.^8^ Cellular (but not plasma) FN is excreted into the urine and expected to get into contact with the apical/ciliary surface of tubular epithelial cells.^9^ To learn more about the specific role(s) of FN in the kidney we generated and investigated Fn-deficient mice. Here we report that the conditional deletion of FN in newborn animals causes loosening of the kidney ECM at its cortico-medullary junction initiating a process that causes progressive cyst formation leading to a substantial loss of parenchyma at the age of 5-6 weeks. Our results indicate that the kidney ECM plays a critical role in preventing cyst formation and phenotypes as seen in nephronophthisis or polycystic kidney disease.

## Material and Methods

### Mice

For *in vivo* experiments we used tamoxifen inducible CAGG-Cre-ER^™^/Fn^fl/fl^ mice as experimental and Fn^fl/fl^ littermates as control mice. The genetic background was C57BL/6J and both sexes were used for the experiments. To generate experimental mice, animals with two floxed alleles of Fn (kindly supplied by Reinhard Fässler, MPI of Biochemistry, Martinsried, Germany) were crossed with CAGG-Cre-ER^™^ mice, that are heterozygous for transgenic CAGG-Cre-ER^™^. Mice were housed under standardized conditions of 62 % air humidity and 21 °C room temperature. Feeding was *ad libitum*. Animals were kept at 12 h light/dark cycle (6 a.m. to 6 p.m.). Genomic DNA from ear biopsies were isolated and tested by PCR. For Cre PCR analysis the primers were as follows: 5’-atgcttctgtccgtttgccg-’3 (sense) and 5’-cctgttttgcacgttcaccg-‘3 (antisense). The thermal cycle profile was denaturation at 95 °C for 30 sec, annealing at 61 °C for 30 sec and extension at 72 °C for 35 sec for 35 cycles. For FN PCR analysis the primers were as follows: 5’-gccatgatctcacactgtagc-’3 (sense) and 5’-ttgccaactgacttggtgag-’3 (antisense). The thermal cycle profile for FN PCR was denaturation at 95 °C for 30 sec, annealing at 60 °C for 30 sec and extension at 72 °C for 1 min for 35 cycles. All experiments conformed to the tenets of the National Institutes of Health Guidelines on the Care and Use of Animals in Research, the EU Directive 2010/63/E and the Association for Research in Vision and Ophthalmology Statement for the Use of Animals in Ophthalmic in Vision Research and were approved by local authorities (54-2532.1-44/12; Regierung Oberpfalz, Bavaria, Germany).

### Tamoxifen treatment

CAGG-Cre-ER^™^ mice express a tamoxifen-inducible Cre recombinase under an ubiquitously active CAGG promotor. Binding of tamoxifen results in the dissolution of the recombinase-receptor complex of Hsp90, which allows Cre recombinase to enter the nucleus and initiate recombination. Tamoxifen was diluted in corn oil to a final concentration of 2.5 mg/ml and administered via eye drops (10 µl/eye). Both experimental mice and their respective control littermates were equally treated with tamoxifen eye drops three times per day in four-hour intervals as described previously.^10^ Treatment started on postnatal days (P) 1, P5, and P10, respectively, and was applied for either 2 or 5 days. Kidneys were harvested and investigated at P3, P4, P8, P14, P18, and 5-6 weeks of age.

### mT/mG reporter mice

mT/mG reporter mice were used to confirm successful activation of the Cre recombinase via tamoxifen eye drops. mT/mG reporter mice express a membrane-targeted green fluorescent protein after Cre-mediated excision.^11^ CAGG-Cre-ER^™^ mice were crossed with homozygous mT/mG mice. The offspring was treated with tamoxifen eye drops from P1 to P5, according to protocols published previously.^10^ At the age of 5-6 weeks, mice were perfused with 4 % paraformaldehyde (PFA). Kidneys were removed, fixed overnight in 4 % PFA, washed extensively in phosphate buffer (PB, 0.1 mol/L, pH 7.4), incubated in 10 %, 20 % and 30 % sucrose overnight, and shock frozen in tissue-mounting medium (O.C.T Compound; DiaTec, Bamberg, Germany). Frozen sections were washed three times in PB (10 min each), and cell nuclei were counterstained with 4,6-diamidino-2-phenylindoel (DAPI, Vectashield; Vector Laboratories, Burlingame, USA) 1:10 diluted in fluorescent mounting medium (Serva; Heidelberg, Germany).

### Ventricular perfusion

Mice were deeply anesthetized by intraperitoneal injections containing ketamine (120 mg/kg body weight) and xylazine (8 mg/kg body weight). Mice were perfused through the left ventricle with 0.89 % NaCl, followed by 4 % PFA for fixation. Kidney and liver tissue was removed and either further fixed in 4 % PFA overnight for histology and electron microscopy or transferred to peqGold Trifast^™^ (Peqlab Biotechnologie GmbH, Erlangen, Germany) for RNA analysis and to dissecting buffer for protein analysis, respectively.

### RNA analysis

Total RNA from kidney and liver samples was extracted using peqGold Trifast^™^ according to manufacturer’s instruction. The extracted total RNA was subsequently transcribed into first-strand cDNA using the iScript^™^ cDNA Synthesis Kit (Quanta BioSciencies Gaithersburg, USA) according to the manufacturer’s recommendations. Real-time RT-PCR was performed on a BioRad CFX Real-Time PCR Detection System (BioRad, Munich, Germany) using the following temperature profile: 50 cycles of 20 s melting at 94 °C, 10 s of annealing at 60 °C, and 20 s of extension at 60 °C. Primer pairs (**Table 1**) were purchased from Invitrogen (Carlsbad, USA) and extended over exon-intron boundaries. Not reversely transcribed RNA served as a negative control. In initial experiments, we identified *GNB2L* as the most suitable housekeeping gene. Quantification was performed using BioRad CFX Manager software version 3.1.1517.0823 (BioRad).

**Table 1.**
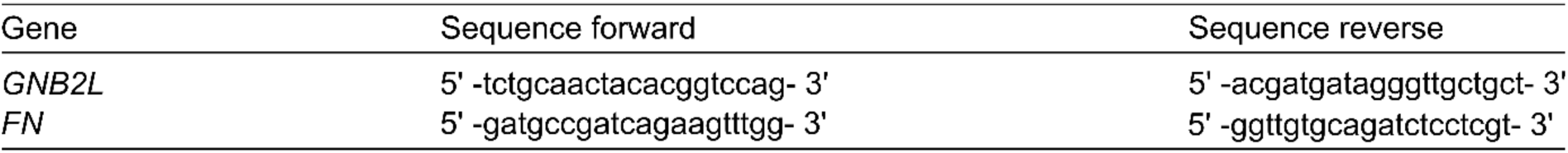
primers used for Real-time PCR

### Western blot analysis

Proteins were isolated in dissecting buffer (0.3 M sucrose, 25 nM imidazole, 0.5 M EDTA) containing protease and phosphatase inhibitors. After a centrifugation step of T5 min at 10000 rpm (4 °C), protein content was measured with the bicinchoninic acid assay (Interchim, Montluçon Cedex, France). For Western blot analysis the proteins were electrophoretically separated on a 10 % polyacrylamide gel and transferred onto a polyvinyldifluoride membrane (PVDF, Millipore). The PVDF membranes were blocked in 5 % nonfat dry milk diluted in Tris-buffered saline with 0.05 % Tween-20 for 1 h at room temperature and then incubated overnight at 4 °C with antibodies against FN (1:70000, rabbit anti-fibronectin, DAKO, Hamburg, Germany). After washing with TBS-T, the PVDF membranes were incubated for 1 h at RT with corresponding secondary antibodies (1:2000, donkey anti-rabbit-horseradish peroxidase, Rockland, Philadelphia, USA) and were subjected to LAS 3000 imaging workstation (Fujifilm, Düsseldorf, Germany), a chemiluminescence detection system. For normalization, whole protein was used as a loading control. The intensity of the bands detected by Western blotting was determined using AIDA Image analyzer software (Raytest, Straubenhardt, Germany).

### Electron and light microscopy

For transmission electron microscopy, kidneys were harvested and investigated at the age of P4, P8 and 5-6 weeks. Kidneys were fixed in 2.5 % glutaraldehyde and 2.5 % PFA in 0.1 M cacodylate buffer for 24 h.^12^ After washing with 0.1 M cacodylate buffer, kidneys were postfixed in 1 % OsO_4_ and 0.8 % potassium ferrocyanide in 0.1 M cacodylate buffer for 2 h at 48 °C. For dehydration kidneys went through a graded series of ethanol followed by Epon (Serva, Heidelberg, Germany) embedding. Semi-thin sections were stained with Richardson’s stain.^13^ Ultrathin sections (60 nm) were mounted on uncoated copper grids, stained with uranyl acetate and lead citrate, and examined on a Zeiss Libra transmission electron microscope (Carl Zeiss AG). For light microscopy kidneys of different ages (P3, P4, P8, P14, P18, 5-6 weeks) were analyzed. After ventricular perfusion, the kidneys were fixed in 4 % PFA overnight, washed extensively in PB (0.1 mol/L, pH 7.4) and equilibrated in 50 % isopropanol and 70 % isopropanol. Afterwards, kidneys were embedded in paraffin, according to standard protocols. Paraffin sections (6 µm) were deparaffinized and washed with water. The sections were stained with hematoxylin and eosin (HE) or with Alcian blue and analyzed using a Zeiss Axio Imager microscope (Carl Zeiss AG, Jena, Germany).

### Immunohistochemistry

Tissue samples were fixed in 4 % PFA overnight, washed in PB (0.1 mol/L, pH 7.4) and embedded in either Tissue-Tek optimal cooling temperature compound (Sakura Finetek Europe B.V., Zoeterwounde, Netherlands) for frozen sections or paraffin. Frozen kidney sections (12 µm) were cut on a cryostat. For detection of CD31 sections were blocked with 2 % bovine serum albumin (BSA), 0.2 % cold water fish gelatin (CWFG), 0.1 % Triton-X-100 in PB (0.1 mol/L, pH 7.4) and incubated with goat anti-CD31 (1:100, R&D systems, Minneapolis, USA). As secondary antibody biotinylated anti-goat IgG (1:500, Vector Laboratories, Burlingame, USA) was used and incubated with Alexa Fluor® 488 streptavidin (1:1000, Life Technologies, Paisley, England). Paraffin sections (6 µm) were deparaffinized and washed with water. For calbindin, megalin and aquaporin-2 staining, sections were pretreated with boiled Tris-EDTA for 40 min. Afterwards, the sections were blocked with 1 % BSA in phosphate-buffered saline (PBS, Invitrogen, Karlsruhe, Germany) for 1 h at room temperature. After blocking, sections were incubated with mouse anti-calbindin (1:400, Swant, Marly, Swiss), goat anti-megalin (1:200, Santa Cruz Biotechnology, Santa Cruz, USA) or goat anti-aquaporin-2 (1:200, Santa Cruz Biotechnology, Santa Cruz, USA). Goat anti-mouse IgG (1:2500, Life Technologies, Paisley, England) conjugated to Alexa Fluor® 488 or donkey anti-goat IgG (1:2500, Jackson Immunoresearch, Baltimore, USA) conjugated to Cy3 were used as secondary antibodies. For fibronectin, collagen IV and laminin staining, the sections were pretreated with Tris-HCl and proteinase K (20 mg/mL, Sigma-Aldrich, Taufkirchen, Germany). Afterwards, the sections were blocked with 2 % BSA in PB (0.1 mol/L, pH 7.4) for 1 h at room temperature. After blocking, sections were incubated with rabbit anti-fibronectin (1:500, DAKO, Hamburg, Germany), rabbit anti-collagen IV (1:200, Rockland, Pennsylvania, USA) or rabbit anti-laminin (1:200, DAKO, Hamburg, Germany). As secondary antibody goat anti-rabbit IgG (1:2500, Jackson Immunoresearch, Baltimore, USA) conjugated to Cy3 was used. For PDGFR-β staining, the sections were blocked with 1 % BSA in PBS for 1 h at room temperature and incubated with rabbit anti-PDGFR-β (1:200, Abcam, Cambridge, England) overnight. Goat anti-rabbit IgG (1:2500, Jackson Immunoresearch, Baltimore, USA) conjugated to Cy3 was used as secondary antibody. For hyaluronan staining, the sections were blocked with 2 % BSA in PB (0.1 mol/L, pH 7.4) and incubated with rabbit anti-hyaluronan binding protein (1:50, amsbio, Abingdon, England) conjugated to biotin (B-HABP). As secondary antibody Alexa Fluor® streptavidin (1:1000, Life Technologies, Paisley, England) was used. Primary antibodies were always incubated overnight at 4 °C and secondary antibodies were incubated for 1 h at room temperature in the dark. Finally, DAPI (Vector Laboratories) was added to counterstain nuclear DNA. Slides were analysed using a Zeiss Axio Imager microscope (Carl Zeiss AG, Jena, Germany).

### RNAscope In Situ Hybridization

Kidneys were obtained from Fn^fl/fl^ control animals at the age of P4 and fixed overnight in 4 % PFA. Next, kidneys were washed extensively with PB (0.1 mol/L, pH 7.4) and after a series of isopropanol embedded in paraffin. Kidney sections (5 µm) were deparaffinized and *in situ* hybridization was performed according to the manufacturer’s instructions.^14^ To detect mRNA, the RNAscope Multiplex Fluorescent v2 kit (Advanced Cell Diagnostics ACD, Hayward, USA) was used. For Fn mRNA detection, the RNAscope probe-Mm-Fn1 (Advanced Cell Diagnostics ACD, Hayward, USA) was used. Additionally, Opal fluorophores 690 (Akoya Biosciences, Marlborough, USA) in tyramide signal amplification buffer was used to detect hybridization signals. Finally, nuclei were counterstained with DAPI included in the Multiplex Fluorescent v2 kit. Slides were analysed using a Zeiss Axio Imager microscope (Carl Zeiss AG, Jena, Germany).

## Results

### Generation of newborn mice with induced FN deficiency

To learn about the importance and specific role(s) of FN in the kidney, we established a mouse model with an ubiquitous deficiency of FN using the tamoxifen-inducible Cre/loxP system. Since the general deletion of FN in the mouse embryo leads to early embryonic lethality before embryonic day (E) 10.5,^15^ we aimed at deleting FN after birth. To this end we obtained Fn^fl/fl^ mice^16^ (kindly supplied by Reinhard Fässler, MPI of Biochemistry, Martinsried, Germany) with floxed alleles of Fn. Fn^fl/fl^ mice were crossed with tamoxifen-dependent CAGG-Cre-ER^™^ transgenic mice. To initiate FN deficiency in newborn mouse pups we used a non-invasive protocol using tamoxifen eye drops that was developed in our laboratory.^17^ In our hands, this treatment reliably induces Cre not only in the eye, but also in tissues throughout the body including the kidney, presumably due to its metabolization by corneal CYP2D6 (a cytochrome P450 isoform) and uptake from the ocular surface into the general circulation.^18, 19^ Successful induction was first demonstrated by crossing CAGG-Cre-ER^™^ mice with mT/mG mice, a double-fluorescent Cre reporter mouse strain that expresses membrane-targeted green fluorescent protein (mG) after excision, reflecting activation of the Cre recombinase.^11^ To this end the experimental animals were treated from P1 to P5 three times daily with tamoxifen eye drops (2.5 mg/ml) and the isolated tissue was examined for fluorescence at 5-6 weeks of age. Littermates that were also treated with tamoxifen but did not express Cre recombinase functioned as a control group (Fn^fl/fl^). Fluorescence microscopy of frontal sections of the kidney showed a specific GFP signal in the glomeruli, tubules and blood vessels in CAGG-Cre-ER^™^/mT/mG animals, indicating successful induction of Cre recombinase via eye drops (**Figure 1A**). No expression of GFP was detectable in the control animals (**Figure 1A**).

**Figure 1.**
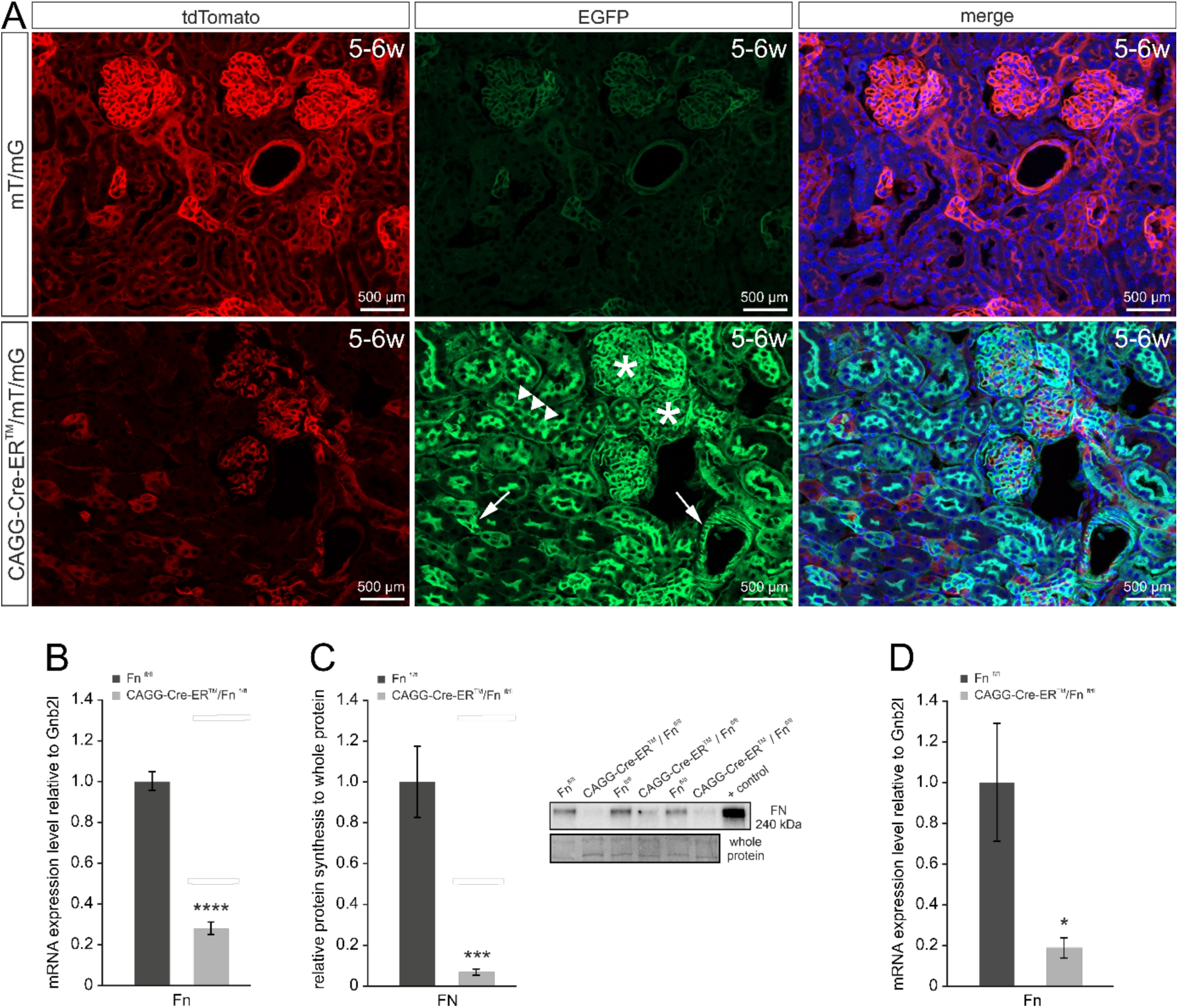
Induction of FN deficiency in the kidney. **A**. Successful Cre-recombinase mediated recombination in kidneys of mT/mG mice after tamoxifen treatment. For induction, mice were treated from P1 to P5 with tamoxifen eye drops (2.5 mg/ml) three times a day. Fluorescence microscopy analysis of frontal sections of 5-6-week-old CAGG-Cre-ER^™^/mT/mG kidneys show specific GFP staining in the glomeruli (asterisk), tubules (triangle) and blood vessels (arrow) reflecting successful excision of the membrane-targeted green fluorescent protein dimer Tomato; no corresponding signal is detectable in control mT/mG mice. **B**. Significant reduction of FN in kidneys of CAGG-Cre-ER^™^/Fn^fl/fl^ mice 4 days after the start of induction. For induction, mice were treated from P1 to P2 three times daily with tamoxifen eye drops (2.5 mg/ml). Isolation of renal mRNA and proteins was performed at P4. Quantitative real-time RT-PCR analyses show a highly significant reduction of renal Fn mRNA expression in CAGG-Cre-ER^™^/Fn^fl/fl^ animals compared to control animals (n = 4-7; mean ± SEM; ****p < 0.001). **C**. Western blot analysis and the corresponding relative densitometric analysis for FN in kidneys confirm a significant decrease of FN in the CAGG-Cre-ER^™^/Fn^fl/fl^ mice at the age of P4 compared to the controls (n = 6; mean ± SEM; ***p < 0.005). Whole protein was used as loading control for normalization and recombinant FN as positive control. **D**. Significant reduction of Fn mRNA in livers of CAGG-Cre-ER^™^/Fn^fl/fl^ mice. For induction, mice were treated from P1 to P5 three times daily with tamoxifen eye drops (2.5 mg/ml). Isolation of hepatic mRNA was performed at the age of 5-6 weeks. Quantitative real-time RT-PCR analyses show a highly significant reduction of Fn mRNA expression in livers of CAGG-Cre-ER^™^/Fn^fl/fl^ animals compared to control animals (n = 4; mean ± SEM; *p < 0.05).

In a next step CAGG-Cre-ER^T^/ Fn^fl/fl^ animals were treated with tamoxifen eye drops from P1 to P2 three times daily. Already two days later at P4, quantitative real-time RT-PCR showed a highly significant reduction (p < 0.001) in the amounts of mRNA in CAGG-Cre-ER^T^/Fn^fl/fl^ kidneys compared to the control Fn^fl/fl^ littermates (**Figure 1B**). Densitometric evaluation of Western blot assays (p < 0.005) with proteins isolated from kidneys also indicated a highly significant reduction in renal FN expression in CAGG-Cre-ER^™^/Fn^fl/fl^ mice compared to control animals (**Figure 1C**). Since plasma-derived FN is incorporated in the extracellular matrix of several organs including the kidney and FN is produced and secreted by hepatocytes ^8^ we verified if tamoxifen treatment also altered the mRNA expression of Fn in the liver. Quantitative real-time RT-PCR of 5-6 weeks old CAGG-Cre-ER^™^/Fn^fl/fl^ livers, treated with tamoxifen eye drops from P1 to P5 showed a significant reduction in the amounts of Fn mRNA of approximately 80 % in comparison to the control Fn^fl/fl^ animals (**Figure 1D**).

### Cyst formation in FN-deficient mice

Induction of ubiquitous FN deficiency in newborn CAGG-Cre-ER^™^/Fn^fl/fl^ mice with tamoxifen eye drops led to the formation of renal cysts that were filled with clear fluid or blood and replaced almost completely intact parenchyma at 5-6 weeks of age (**Figure 2A**). No obvious structural changes were observed in other organs including liver, lung, spleen, heart, pancreas, salivatory glands, eye or brain (data not shown). Next we treated mice with tamoxifen eye drops three times a day starting from P1 to obtain information on the time course of cyst formation in the kidney. Kidneys were histologically examined by HE staining 3, 4, 8 or 14 days after the start of induction. Here, animals whose renal changes were examined at P3 and P4 received a 2-day tamoxifen treatment, while the induction period for the remaining experimental animals extended to a period of 5 days. The histological analysis showed that on P3 no pathological changes were visible in CAGG-Cre-ER^™^/Fn^fl/fl^ kidneys in comparison to the controls (**Figure 2B**). In contrast 24 hours later (P4) first cysts were observed in animals with a deletion of FN. In P4 kidneys the cysts were mainly localized in the transition area between cortex and medulla. With increasing age, the cysts enlarged. Already on P8, the cysts were displacing intact kidney tissue (**Figure 2B**).

**Figure 2.**
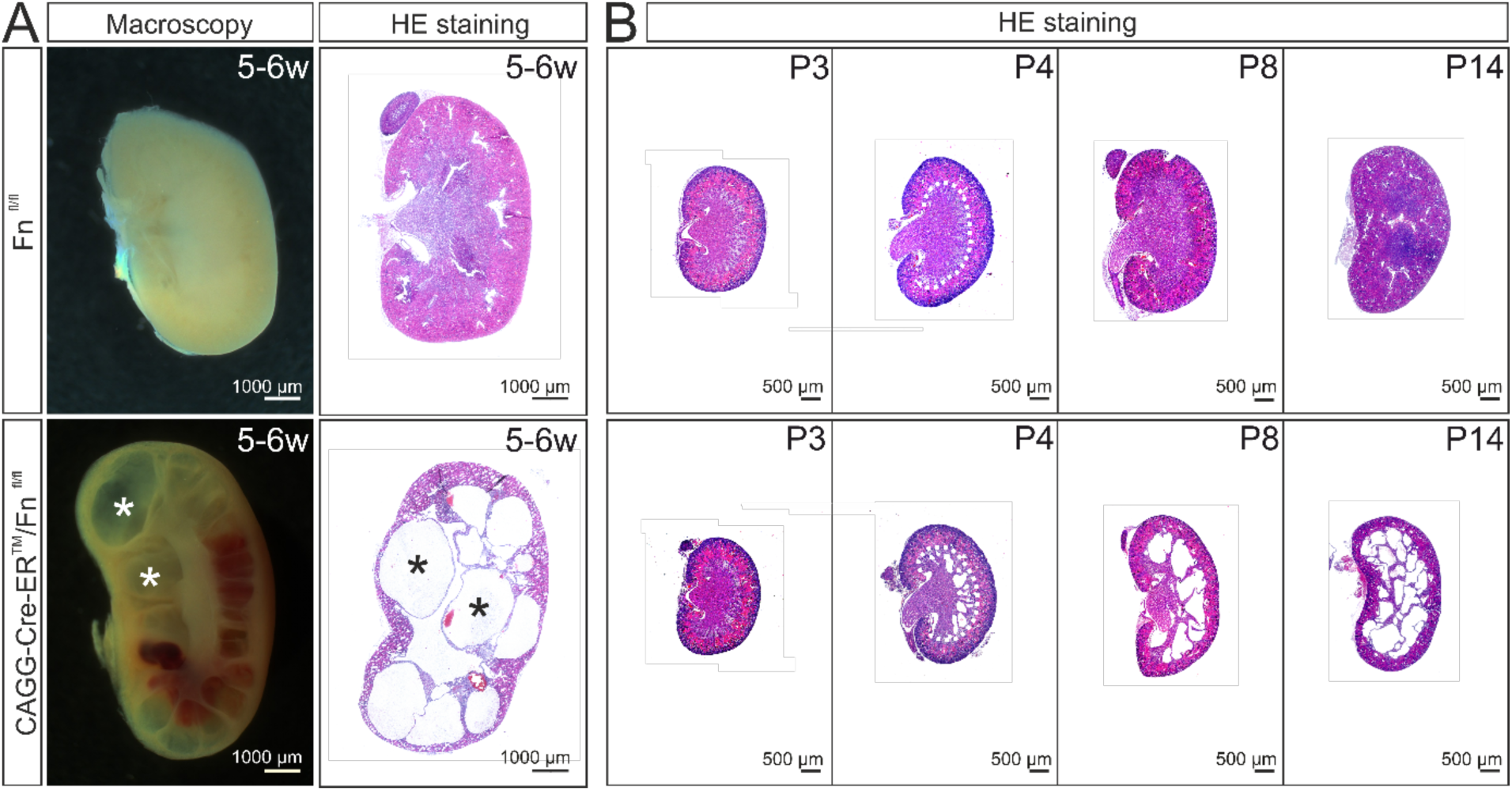
Cyst formation in CAGG-Cre-ER^™^/Fn^fl/fl^ mice after tamoxifen-induced deletion of FN. **A**. Macroscopic images (left), as well as histological staining of the kidneys (right) at the age of 5-6 weeks document the presence of numerous fluid-filled cyst (asterisk) in CAGG-Cre-ER^™^/Fn^fl/fl^ mice. For induction of FN deficiency, animals were treated from P1 to P5 with tamoxifen eye drops (2.5 mg/ml) three times a day. **B**. Progressive course of cyst formation in CAGG-Cre-ER^™^/Fn^fl/fl^ mice after tamoxifen-induced deletion of FN. For induction, the experimental animals were treated with tamoxifen eye drops (2.5 mg/ml) three times a day from P1. Histological analyses of the kidneys were performed at 3, 4, 8, 14 days and 5-6 weeks of age, respectively. Representative HE stains indicate a positive correlation between the age of the animals and the size and/or number of cysts. First cysts appear at the age of P4 at the transition area between medulla and cortex (dotted line).

### *Localization of* Fn *expression in the kidney*

Based on the quite specific localization of the cysts at the medulla-cortex boundary we wondered if this is correlated with the presence of FN protein in this area. Indeed, we found a strong interstitial, radial-orientated expression pattern of FN protein lining the nephrons between cortex and medulla when P4 Fn^fl/fl^ kidneys were stained with antibodies against FN (**Figure 3A**). Positive signal was also detectable in the cortical interstitium and the mesangial ECM of the glomeruli, but to a lesser extent (**Figure 3A**). Immunohistochemical stainings of CAGG-Cre-ER^™^/Fn^fl/fl^ kidneys at P4 showed reduced amounts of protein throughout the kidney further confirming the successful deletion of Fn (**Figure 3A**). In addition, we performed RNAscope to detect Fn mRNA expression in Fn^fl/fl^ and CAGG-Cre-ER^™^/Fn^fl/fl^ kidneys. At P4 the signal for Fn mRNA expression in control Fn^fl/fl^ kidneys strongly corresponded to the location of protein expression (**Figure 3A**). Accordingly it was found predominantly in the outer medulla (**Figure 3B**). *In situ* hybridization of kidneys harvested at P14 and 5-6 weeks showed a shift of mRNA expression from the interstitial tissue of the outer medullary area to the cortex (**Figure 3B**). While P14 kidneys still expressed Fn mRNA in the outermost part of the medulla, in 5-6 weeks old kidneys the expression was almost completely restricted to the cortex (**Figure 3B**).

**Figure 3.**
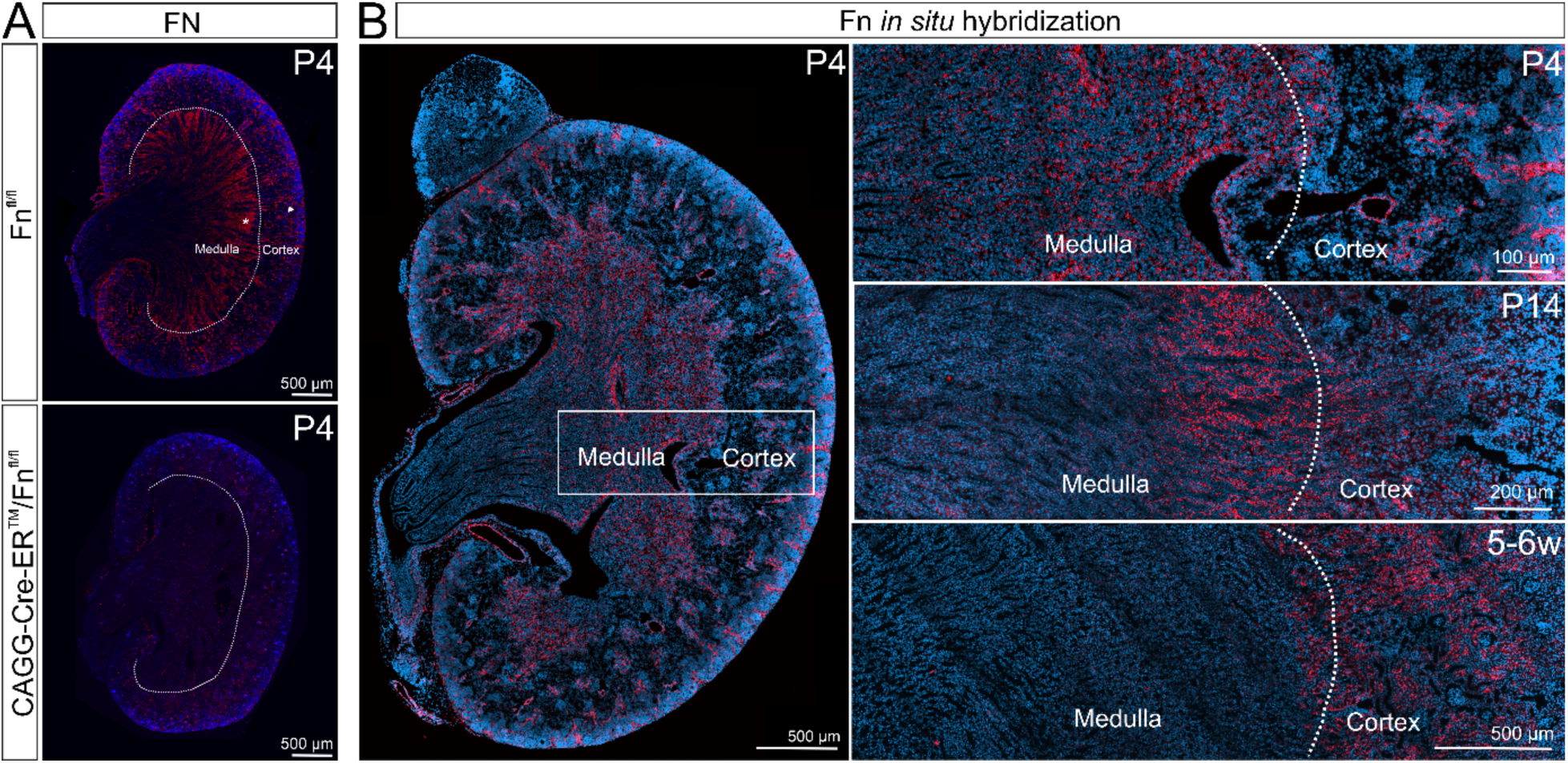
Localization of Fn mRNA expression strongly correlates with FN protein expression and relocates over time. For induction, the mice were treated from P1 to P2 three times daily with tamoxifen eye drops (2.5 mg/ml). **A**. Fluorescence microscopy analysis of frontal sections of whole Fn^fl/fl^ kidneys at P4 shows positive interstitial (asteriks) and mesangial (triangle) staining throughout the organ with high amounts of FN protein in the outer medulla close to the medulla-cortex boundary (dotted line). In contrast, CAGG-Cre-ER^™^/Fn^fl/fl^ mice display reduced amounts of FN protein. Red = FN, blue = DAPI. **B**. Localization of Fn mRNA expression in Fn^fl/fl^ kidneys at P4 by *in situ* hybridization using RNA scope strongly correlates with the protein expression seen in **A** (whole kidney section left and higher magnification top). Again, highest level of FN expression is found in the medullary interstitium near the medulla-cortex border (**B**, higher magnification top). In addition, expression is present in the cortical interstitium and the glomeruli. Higher magnifications (**B**, middle and bottom) show Fn^fl/fl^ kidneys at the age of P14 and 5-6 weeks. A clear relocation of Fn mRNA is visible from the outer medulla to the cortex. At the age of 5-6 weeks Fn mRNA is almost exclusively expressed in the cortex (**B**, bottom). Red = Fn mRNA, blue = DAPI

### Cellular origin of cysts in FN-deficient mice

Since there is evidence that PKD cysts derive from nephrons and collecting ducts,^20^ we stained P4, P8 and 5-6-week-old kidneys for calbindin as a marker for distal convoluted tubules (**Figure 4A**), for megalin as a marker for proximal convoluted tubules (**Figure 4B**), and for aquaporin-2 which is a marker for collecting ducts (**Figure 4C**). When we stained for calbindin and megalin we found a specific positive signal, as expected, in distal and proximal convoluted tubules of both Fn^fl/fl^ and CAGG-Cre-ER^™^/Fn^fl/fl^ kidney sections but no positive immunoreactivity in the cells located adjacent to the cyst (**Figure 4A/B**). The same result was obtained when we stained for aquaporin-2. We found positive immunoreactivity in collecting ducts but no signal in cells along the cysts (**Figure 4C**). Next, we wondered if the cysts might be dilated blood vessels. Still, staining with antibodies against CD31 as a marker for endothelial cells showed, as expected, in both, Fn^fl/fl^ and CAGG-Cre-ER^™^/Fn^fl/fl^ kidneys positive immunoreactivity in cells of peritubular capillaries (**Figure 4D**) but no signal in the cells lining the cysts (**Figure 4D**).

**Figure 4.**
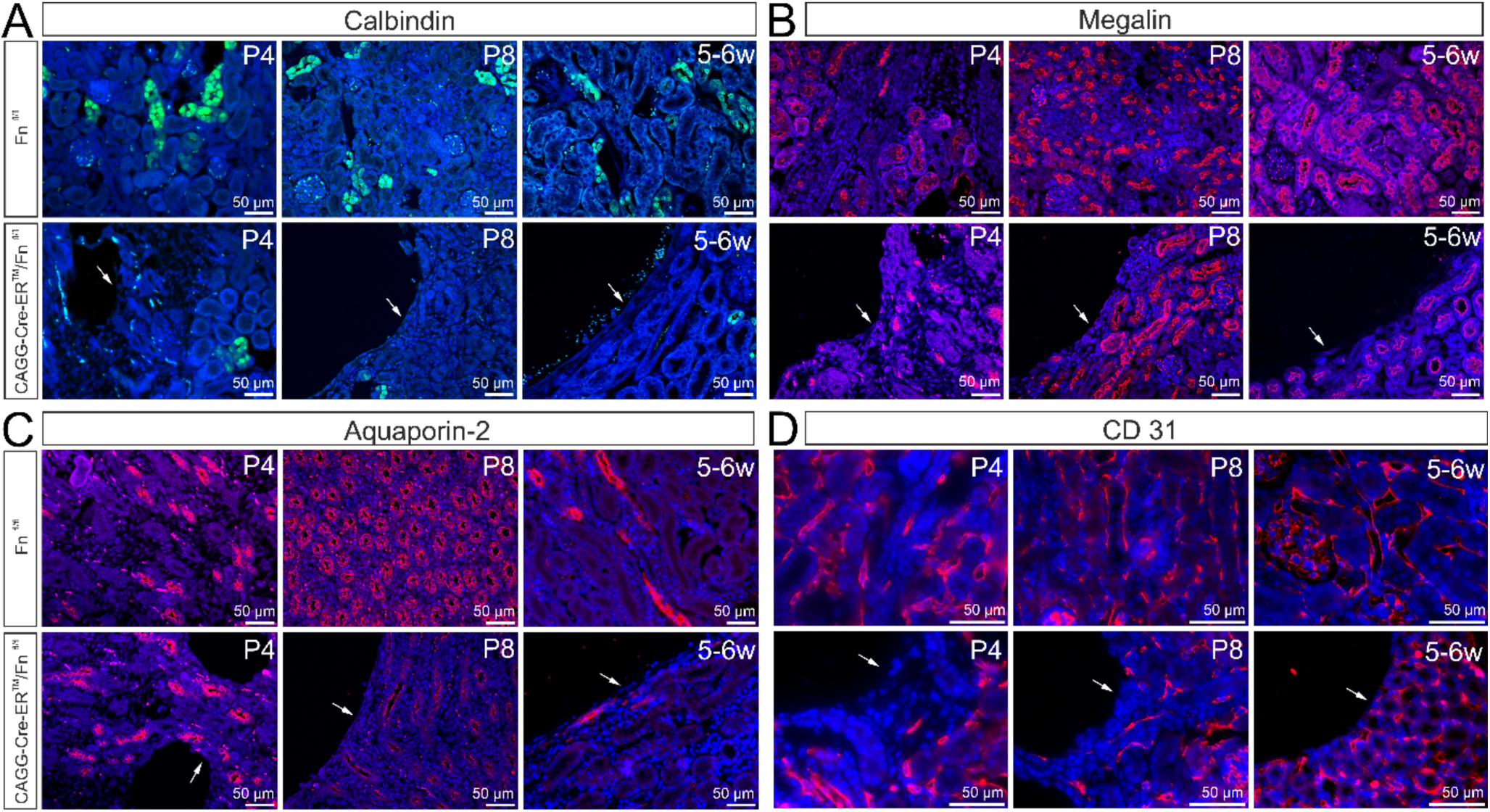
Cellular origin of cysts in FN-deficient kidneys. For induction, the experimental animals were treated with tamoxifen eye drops (2.5 mg/ml) three times a day from P1. Immunohistochemical analyses of the kidneys were performed at 4, 8 days and 5-6 weeks of age, respectively. **A**. Immunohistochemistry with antibodies against calbindin shows representative staining in distal convoluted tubules in both, CAGG-Cre-ER^™^/Fn^fl/fl^ and control mice, but no positive signal in cells lining the cysts (bottom row, arrow). Green = calbindin, blue = DAPI. **B**. Staining of megalin shows representative staining in proximal convoluted tubules in both, CAGG-Cre-ER^™^/Fn^fl/fl^ and control mice, but no positive signal in cells lining the cysts (bottom row, arrow). Red = megalin, blue = DAPI. **C**. Staining of aquaporin-2 shows, as expected, immunoreactivity in collecting ducts in the experimental and control group, but again no staining in cells along the cysts (arrow) in CAGG-Cre-ER^™^/Fn^fl/fl^ animals. Red = aquaporin-2, blue = DAPI. **D**. Staining of kidneys with antibodies against CD31 could exclude endothelial cells as cellular source of cyst formation because none of the cells lining the cysts (arrow) are positive for CD31. Endothelial cells of peritubular capillaries are positive for CD31 in CAGG-Cre-ER^™^/Fn^fl/fl^ and Fn^fl/fl^ control littermates. Red = CD31, blue = DAPI.

To characterize the changes in fine structure that are associated with cyst formation after FN deletion, semi-thin sections were stained with Richardson’s stain and examined in detail.

Light microscopical investigation of P4 and P8 kidneys showed a clear loosening of the interstitial tissue along the cysts. While on P4 both interstitial cells and tubular structures were located along the cyst (**Figure 5A**), on P8 only interstitial cells were apparent between the cysts and normal kidney parenchyma (**Figure 5A**). At the age of 5-6 weeks, the cysts were surrounded by a membrane-like structure, which suggested a clear demarcation from still intact tissue (**Figure 5A**). By transmission electron microscopy of P4 kidneys, a loose arrangement of cells along the developing cysts was seen (**Figure 5B**). On P8 or 5-6 weeks after the start of induction, the cells along the cyst were flattened, elongated and arranged in an epithelial-like manner (**Figure 5B**). In addition, cell-cell contacts between individual cells were seen (**Figure 5C**). Moreover, there was an increased amount of 5-6 nm thick actin filaments along the cyst boundary (**Figure 5C**).

**Figure 5.**
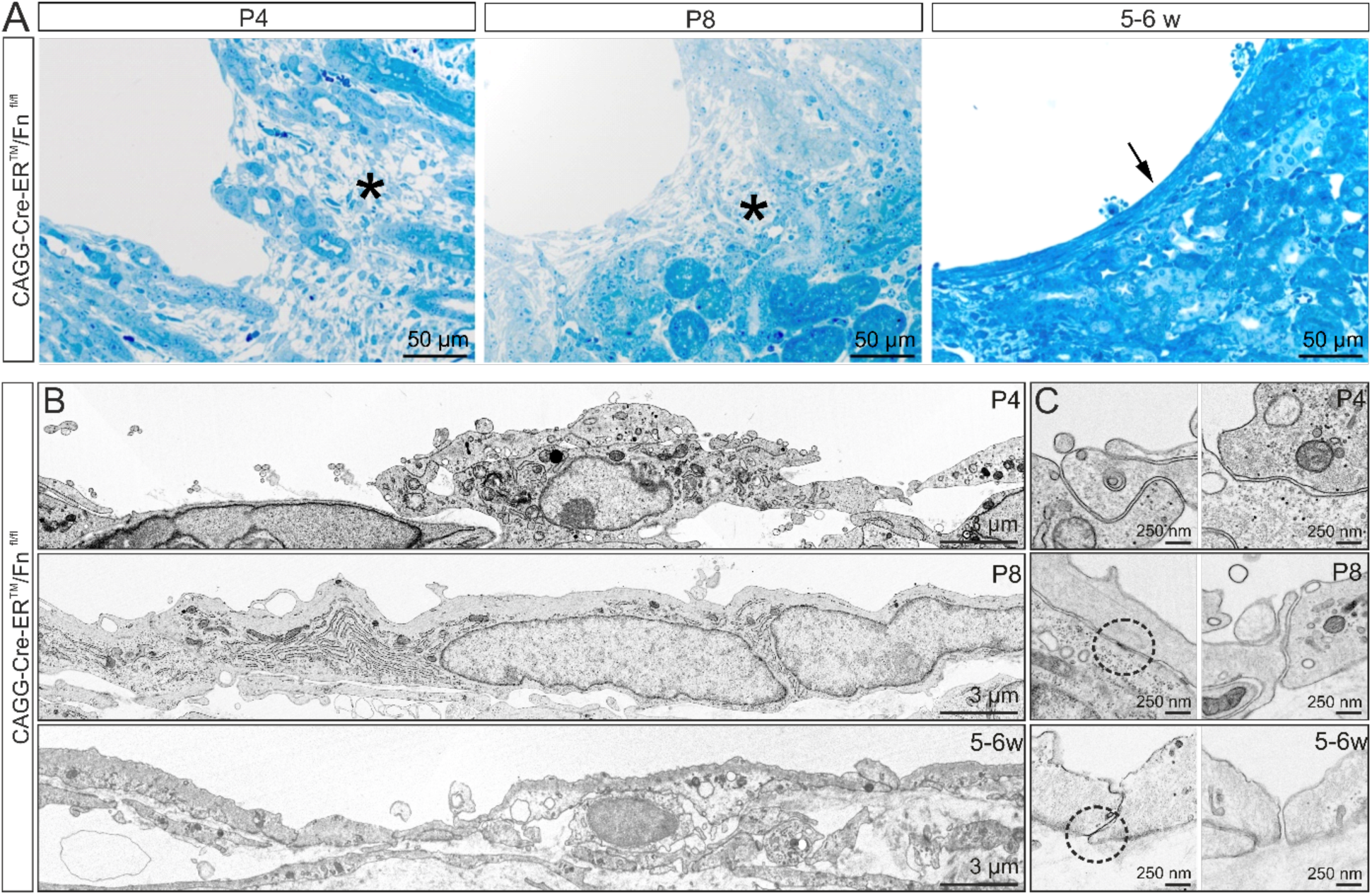
Fine structure of forming cysts in FN-deficient kidneys. For induction, the animals were treated with tamoxifen eye drops (2.5 mg/ml) three times a day from P1. The analyses were performed at 4 or 8 days and 5-6 weeks of age. **A**. Semi-thin sections of cystic kidneys (Richardson`s stain) show a loosening of the interstitial tissue (asterisk) 4 or 8 days after the start of induction, while after 5-6 weeks the cysts appear to be enclosed by an epithelium or epithelial-like layer (arrow). **B**. Electron micrographs of the cyst boundary. The cells along the cyst do not show any tissue-specific morphological features. **C**. In contrast to P4, at P8 or 5-6 weeks after the start of induction, cell contacts between cells are visible (dashed circles).

### Cysts in FN-deficient animals are not lined by an epithelium and are not continuous with lumina of tubules

An epithelial nature of the cells would argue for a tubular origin of the cysts (**Figure 5B and C**). To this end, immunostaining for collagen IV and laminin was performed to check for the presence of a basal lamina. In addition, immunohistochemical staining against E-cadherin was performed to test for epithelial cell-cell contacts. Neither on P4 or P8, nor 5-6 weeks after the start of induction a continuous positive signal for laminin, collagen IV, and E-cadherin could be observed in the cells lining the cysts (**Figure 6A, B and C**). Furthermore, semi-thin serial sections of renal cysts at P4 confirmed their formation due to dilatation of the interstitium. No connection to an adjacent tubule could be observed (**Supplemental Figure 1**). However, as no epithelial-like demarcation of the intact renal tissue could be observed at this stage of cyst development (**Figure 5**), a corresponding analysis was performed again 14 days after the start of induction. Semi-thin serial sections stained with Richardson’s stain showed that on P14 a clear border of the cyst was visible. Although many renal tubules were noted along the cyst -in addition to interstitial cells -no clear fusion with a tubule or other renal structure was evident at this time of examination (**Supplemental Figure 1**).

**Figure 6.**
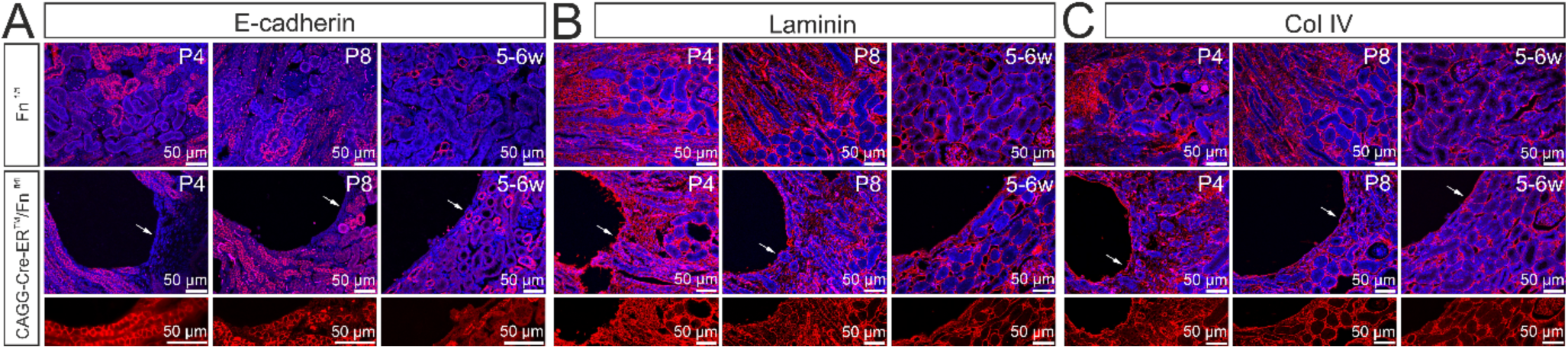
Renal cysts are not lined by an epithelium. For induction, the mice were treated with tamoxifen eye drops (2.5 mg/ml) three times a day starting at P1. The analyses were performed at the age of 4 or 8 days and 5-6 weeks. **A**. Immunohistochemical staining of frontal kidney sections with antibodies against E-cadherin. Both, CAGG-Cre-ER^™^/Fn^fl/fl^ and Fn^fl/fl^ control littermates display the representative signal in junctional contacts of epithelial cells lining distal tubules and collecting ducts. Cell-cell contacts between cells located adjacent to the cysts (arrow) are not detectable. Red = E-cadherin, blue = DAPI. **B**. Immunohistochemistry of frontal sections of the kidney against laminin documents no characteristic staining for a basement membrane along the cysts (arrow). The bottom row shows a section of the cyst boundary. Red = laminin, blue = DAPI. **C**. Immunohistochemical staining of frontal kidney sections against collagen IV documents no basal lamina-specific staining along the cysts (arrow). The bottom row shows a section of the cyst boundary. Red = collagen IV, blue = DAPI.

### Cysts in FN-deficient mice are lined by interstitial cells

Closer examination of HE stained CAGG-Cre-ER^™^/Fn^fl/fl^ kidneys gives the impression that the nephrons are located lateral to the cysts (**Figure 7A**) which let us assume that the cysts might have their origin in the interstitium. Therefore, we used antibodies against PDGFR-β to clarify the role of interstitial cells. Immunohistochemical analysis of PDGFR-β showed expression of the receptor in the interstitial tissue of both groups (**Figure 7B**). Furthermore, in CAGG-Cre-ER^™^/Fn^fl/fl^ mice, cells outlining the cyst were positively stained at P4 and P8, as well as at 5-6 weeks of age (**Figure 7B**).

**Figure 7.**
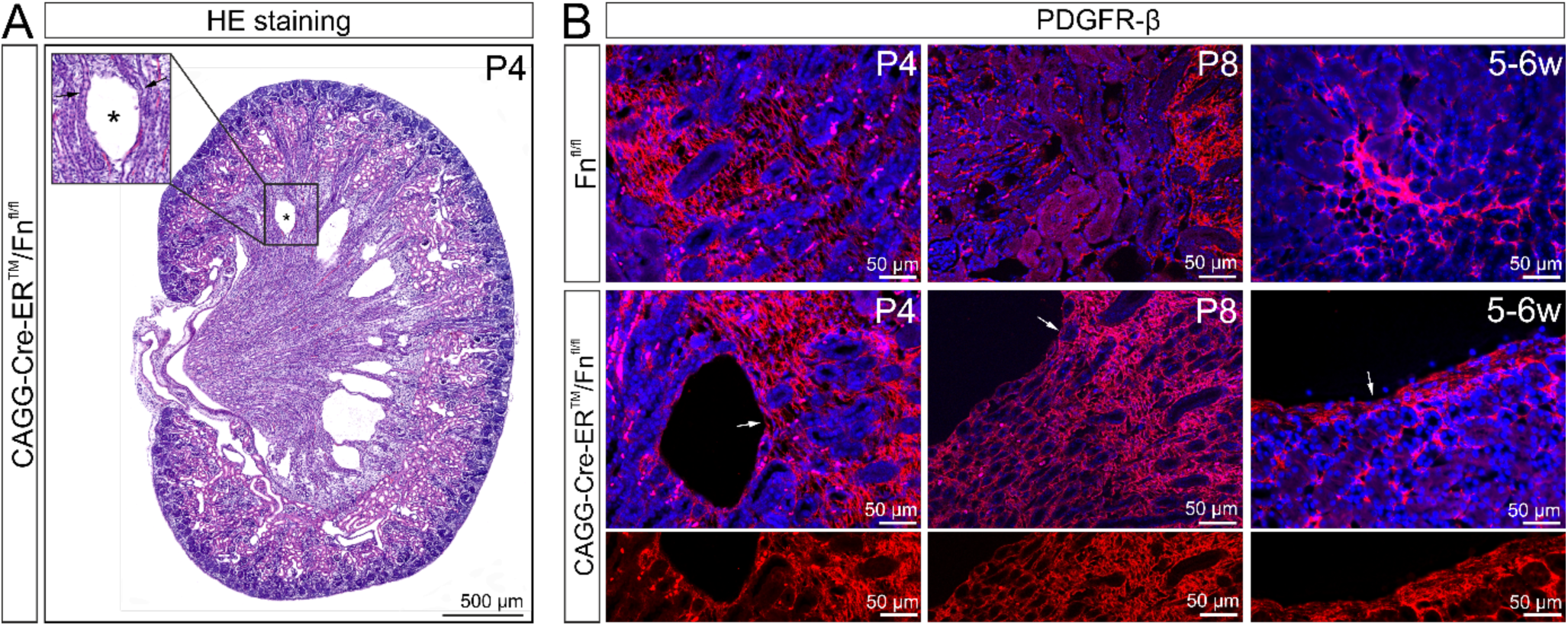
Renal cysts are lined by interstitial cells. For induction, the mice were treated with tamoxifen eye drops (2.5 mg/ml) three times a day starting at P1. The analyses were performed at the age of 4 or 8 days and 5-6 weeks. **A**. Frontal section of a CAGG-Cre-ER^™^/Fn^fl/fl^ kidney at P4 stained with HE with numerous cysts (asterisk) in the outer medulla. Higher magnification (insert) indicates that nephrons (black arrow) are passing the cysts supporting the idea of a loosening process of the interstitium. **B**. Immunohistochemical stainings of frontal kidney sections against PDGFR-β document positive expression along the cyst wall (white arrow) at all observation time points.

### Cysts contain non-fibrillar extracellular matrix

To check if the interstitium is indeed the responsible tissue for the development of renal cysts in FN-deficient mice, non-fibrillar ECM proteins were stained by Hyaluronan-binding protein (HA) and Alcian blue staining. Hyaluronan and proteoglycans that react positive to Alcian blue are essential components of the interstitium ^33^. If the cysts result from the expansion of the interstitium, non-fibrillar components of the ECM should be localized in the cysts. Staining results showed positive reaction for HA and Alcian blue in the renal cysts of CAGG-Cre-ER^™^/Fn^fl/fl^ mice (**Figure 8**).

**Figure 8.**
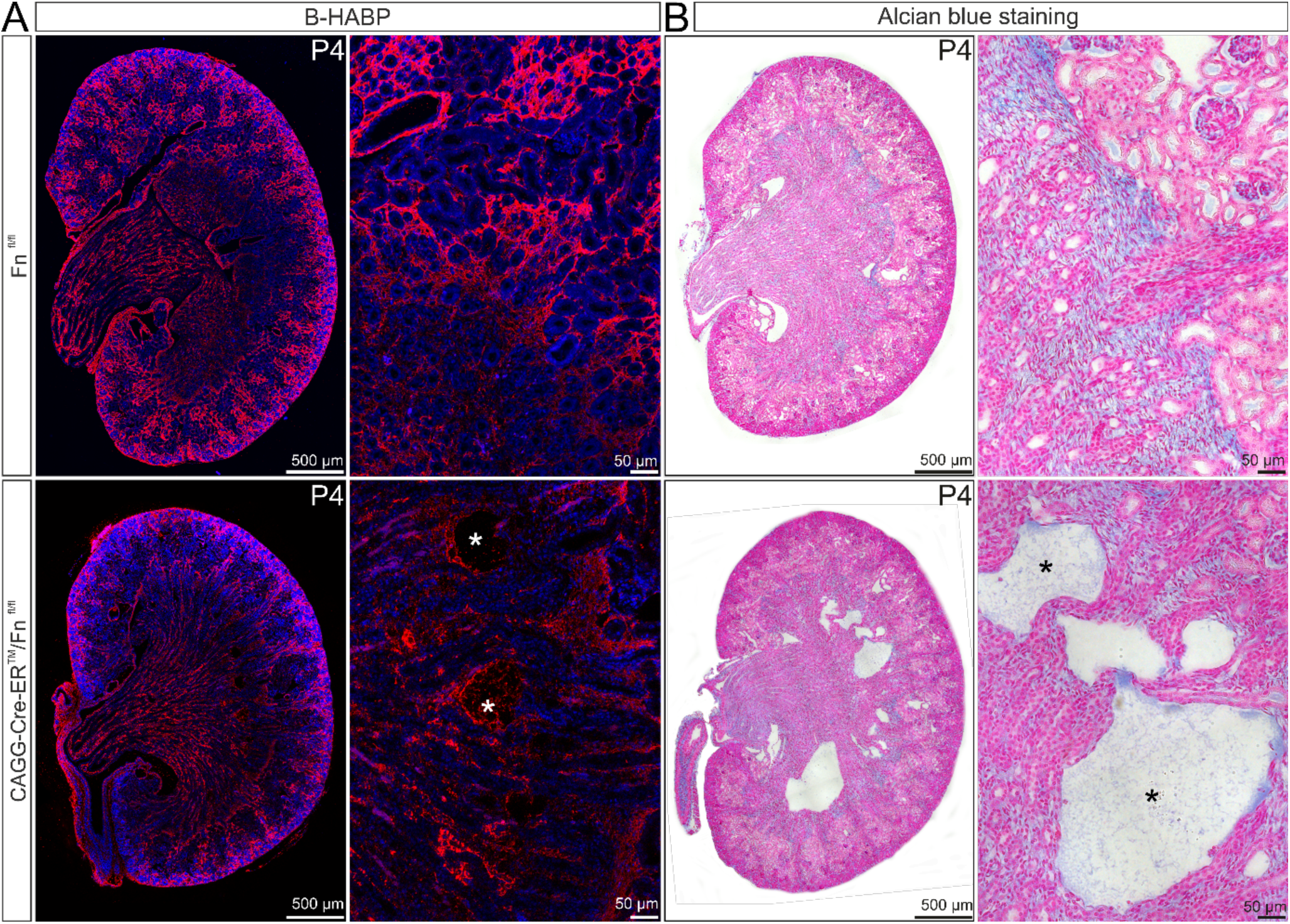
Presence of glycosaminoglycans in renal cysts of CAG-Cre-ER^™^/Fn^fl/fl^ mice. For induction, the mice were treated with tamoxifen eye drops (2.5 mg/ml) three times a day from P1-P2. The analyses were performed on P4. **A and B**. Staining of kidney sections with (**A**) biotinylated hyaluronic acid binding protein (HABP, red) and (**B**) Alcian blue (bluish-green) demonstrate the presence of glycosaminoglycans in the interstitial ECM of both, CAGG-Cre-ER^™^/Fn^fl/fl^ and Fn^fl/fl^ control kidneys (higher magnification in **A** and **B**). Additionally, cysts of FN-deficient kidneys (bottom row) are also filled with positively labelled hyaluronic acid (**A**, white asterisk) and bluish-green dyed glycosaminoglycans (**B**, black asterisk).

### FN-deletion leads to cyst formation at the end of renal development, but not in adult animals

To investigate whether the deletion of FN also leads to renal cyst formation after the end of renal development, mice were treated with tamoxifen eye drops starting at P5 and P10, respectively. HE staining of the kidneys showed that the ubiquitous deletion of FN starting from P5 also leads to the formation of renal cysts (**Figure 9A**). Here, the phenotypic expression of renal cysts was similarly as in CAGG-Cre-ER^™^/Fn^fl/fl^ mice treated with tamoxifen starting at P1 (**Figure 2**). Quantitative real-time RT-PCR assays confirmed a 90% reduction in Fn mRNA levels (p<0.001) in the CAGG-Cre-ER^™^/Fn^fl/fl^ mice (**Figure 9B**). Accordingly, immunohistochemical staining showed a significant reduction in FN in CAGG-Cre-ER^™^/Fn^fl/fl^ mice, in comparison to the control staining (**Figure 9C**). To investigate whether a lack of FN leads to pathological changes between P10 and P14, mice were treated with tamoxifen eye drops three times a day from P10 to P14. HE staining of renal frontal sections showed that deletion of FN beyond P10 does not result in cyst formation (**Figure 9D**). Nevertheless, quantitative real-time RT-PCR analysis revealed a highly significant decrease in Fn mRNA expression of approximately 90 % in the CAGG-Cre-ER^™^/Fn^fl/fl^ mice compared to the controls (p<0.001) (**Figure 9E**). Immunohistochemical staining indicated a moderate reduction of FN compared to control animals (**Figure 9F**).

**Figure 9.**
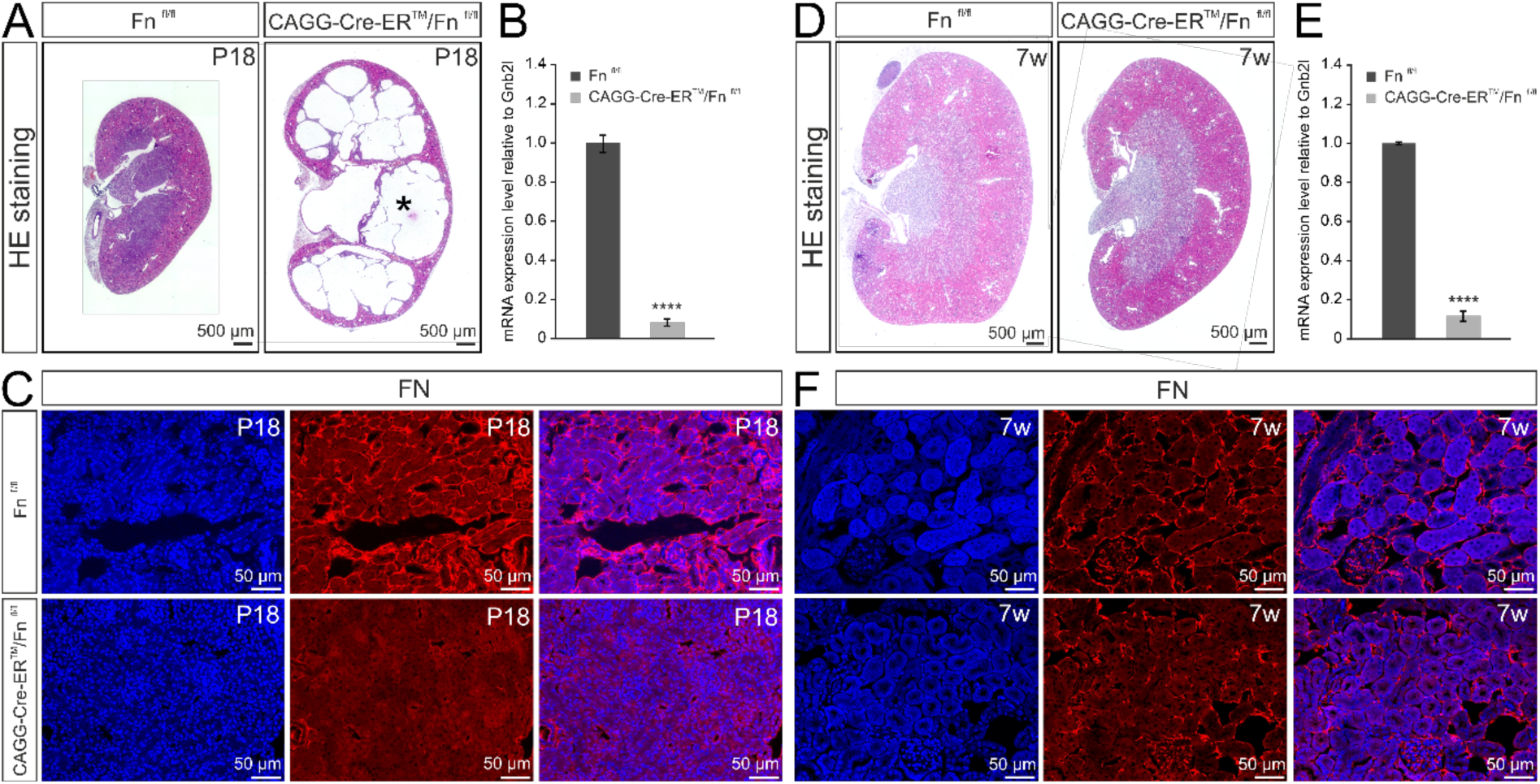
Impact of *Fn* deficiency towards the end of renal development. For induction, the mice were treated from P5 to P9 (**A – C**) and from P10 to P14 (**D – F**), respectively, three times daily with tamoxifen eye drops (2.5mg /ml). The analyses were performed 2 weeks after induction (P5 to P9, **A – D**) or 6 weeks after induction (P10 to P14, **D – F**), respectively. **A - C**. Deletion of Fn from P5 onwards still causes the formation of renal cysts. **A**. HE stained kidney sections of CAGG-Cre-ER^™^/Fn^fl/fl^ and Fn^fl/fl^ control mice. At P18 numerous cysts (asterisk) have formed in CAGG-Cre-ER^™^/Fn^fl/fl^ kidneys. **B**. Quantitative real-time RT-PCR analyses show a highly significant decrease in Fn mRNA expression in CAGG-Cre-ER^™^/Fn^fl/fl^ animals compared to controls (n = 4; mean ± SEM; ****p<0.001). **C**. Immunohistochemical staining against FN on frontal sections of the kidney confirm a clear reduction in the amount of FN in CAGG-Cre-ER^™^/Fn^fl/fl^ mice. **D – F**. Deletion of Fn from P10 onwards does not cause cyst formation. **D**. HE stained kidney sections of CAGG-Cre-ER^™^/Fn^fl/fl^ and Fn^fl/fl^ control mice. 6 weeks after induction, no cysts are visible neither in the Fn^fl/fl^ control mice nor in CAGG-Cre-ER^™^/Fn^fl/fl^ animals. **E**. Quantitative real-time RT-PCR analyses demonstrate a highly significant decrease in Fn mRNA expression in CAGG-Cre-ER^™^/Fn^fl/fl^ mice (n = 3-7; mean ± SEM; ****p<0.001). **F**. Immunohistochemical staining against FN on frontal sections of the kidney indicate a moderate reduction in FN content in CAGG-Cre-ER^™^/Fn^fl/fl^ mice compared to controls.

### Fibrosis in FN-deficient animals

A frequent side effect of kidney damage is the development of fibrosis. This is largely characterized by deposition of collagen fibers.^35^ In order to examine whether the pathological changes associated with the deletion of FN also lead to renal fibrosis, the kidneys were examined for signs of accumulated collagen fibers. Special attention was paid to the tissue adjacent to the cyst. The accumulation of collagen fibers - especially type I and III - was elucidated using Picro-Sirius red staining. At P4, compared to the control group, Picro-Sirius red staining indicated no significant change in the collagen content in the CAGG-Cre-ER^™^/Fn^fl/fl^ mice. There was also no increased collagenous fiber content in vicinity to the developing cyst (**Supplemental Figure 2A**). Histological analysis of animals with a deletion of FN at P8 also did not show any modification in the content of collagen fibers. Similar to P4, the tissue around the cyst was not specifically stained (**Supplemental Figure 2A**). In contrast, at the age of 5-6 weeks, histology showed a clear increase in collagen fibers in the kidneys of CAGG-Cre-ER^™^/Fn^fl/fl^ mice. A striking feature was the increased amount of collagen around the cyst (**Supplemental Figure 2A**). Quantitative real-time RT-PCR analysis of whole kidney samples showed a trend towards an increase in collagen I expression in the CAGG-Cre-ER^™^/Fn^fl/fl^ mice with increasing age, compared to the controls. In contrast, mRNA expression of collagen III did not appear to be affected by the deletion of Fn (n ≥ 4, n ≥ 7; mean ± SEM) (**Supplemental Figure 2B**).

## Discussion

We conclude that FN is critically required for structural maintenance of the kidney and that loss of FN causes a continuous disintegration of the kidney interstitium. The process initiates cyst formation at the cortical-medullary border and continues until the kidney parenchyma is almost completely replaced by cysts. Cysts are surrounded by interstitial cells that form an epithelial-like capsule while cysts continue to enlarge. The conclusion rests upon (1) the generation of mutant newborn mice with conditional deletion of FN throughout the body including the kidney, (2) the observation of cyst formation in newborn mice within two days of FN deletion, (3) the finding based on serial sections that cysts are not connected to lumina of tubuli, (4) the detection of non-fibrillar ECM components within the cysts and (5) the presence of interstitial cells in immediate contact with cysts.

The onset of cyst formation at the cortico-medullary junction is not unlike to the process of cyst formation in nephronophthisis, a tubulo-interstitial, autosomal recessive cystic kidney disease and leading genetic cause of pediatric end-stage renal disease.^21, 22^ Kidneys of affected patients show cortico-medullary cysts and poor cortico-medullary differentiation. Nephronophthisis belongs to the spectrum of disorders termed ciliopathies which result from failure of monocilia or primary cilia function. Primary cilia are immotile, microtubule-based organelles protruding from the surface of nearly every mammalian cell. Cilia act as antennae-like structures to sense chemical or mechanical extracellular signals transmitted by growth factors, hormones, odorants, developmental morphogens or mechanical cues.^23^ By sensing extracellular signals, primary cilia collect, synthesize, and transmit information about the extracellular space into the cell body to promote fundamental cellular responses such as differentiation, maintenance, growth and proliferation. To serve their function primary cilia are associated with numerous pathways that form complex signaling networks. Important factors that govern those signaling pathways include mitogen-activated protein kinase (MAPK), phosphoinositide 3-kinase (PI3K)-AKT, Hippo, nuclear factor κ light-chain enhancer of activated B cells (NF-κB), mammalian target of rapamycin (mTOR), WNT/β-catenin, WNT/PCP (planar cell polarity), sonic hedgehog and transforming growth factor-β.^24^ As of now, our knowledge on the nature of available ligands to initiate signaling on primary cilia is incomplete. There is evidence though indicating that ECM molecules are capable of binding to integrins that are present on the surface of primary cilia and/or on membrane domains of cilia-associated proteins.^25, 26^ Moreover, several experiments point towards a contributing role of an ECM-integrin interaction in renal cyst formation. Accordingly, the conditional inactivation of integrin β1 in mouse collecting ducts results in a dramatic inhibition of polycystin-1-dependent cystogenesis.^27^ In cases of nephronophthisis, changes in the expression of β integrins were noted.^28^ Moreover, mice carrying a hypomorphic allele of the *Lamα5* gene coding for Laminin α5, a typical component of renal basal laminae, develop PKD.^29^

It is of interest to note that loss of Fn mRNA in one-week-old animals does neither lead to cyst formation nor to a substantial reduction in the amounts of deposited fibrillar FN. We assume that the action of crosslinking extracellular enzymes such as transglutaminases substantially extends the half-life of FN in older animals and delays the process of cysts formation.

*In vitro* studies show that binding of FN to β1 integrins, which are localized on primary cilia, augments cilia-dependent Ca^2+^ signals.^26^ In addition, polycystin-1, a cilia-associated protein that is mutated in autosomal-dominant PKD, binds to FN via its extracellular domains.^30^ In the kidney, cilia project from the epithelial cell surface into the tubular lumen and act as flow sensors as well as signaling centers for inter- and intracellular communication. It is generally assumed that cyst formation in ciliopathies is triggered by an impaired function of the cilia on renal epithelial cells. Still, monocilia are also present on cells of mesodermal origin during development and have been observed in interstitial cells of the testis.^31, 32, 34^ It is tempting to speculate that loss of FN function on monocilia of interstitial cells contributes to cyst formation in FN-deficient mice, a hypothesis that we will address in future studies.

## Data sharing statement

Original data are available from the authors upon reasonable request.

## Acknowledgment

We thank Margit Schimmel, Elke Stauber, Angelika Pach and Silvia Babl for excellent technical support. We thank Reinhard Fässler, who kindly supplied Fn^fl/fl^ mice. This study was supported by a grant from the DFG (SFB 1350, Project B3).

## Supplementary Information

**Supplemental Figure 1.**
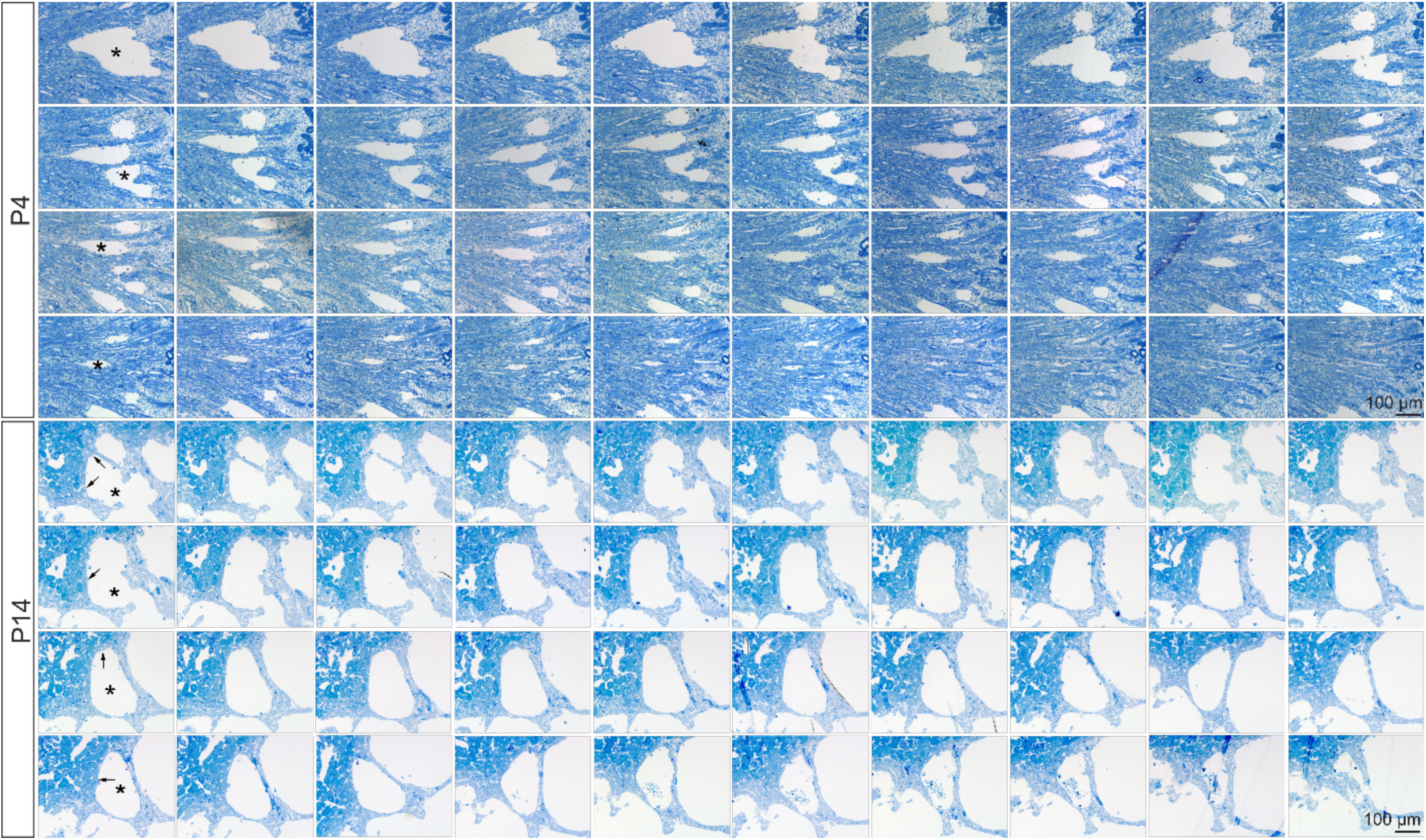
Tubule-independent cystogenesis in FN-deficient mice at P4 and P14. Semi-thin serial sections of a renal cyst at P4 and P14. For induction, the P4 animals were treated three times daily with tamoxifen eye drops (2.5 mg/ml) on P1 and P2. P14 animals were treated three times daily with tamoxifen eye drops (2.5 mg/ml) starting from P1. Histological analysis of the kidney was performed at P4 or P14 by Richardson staining. Serial sections at P4 do not show a transition of the cyst (asterisk) to a renal structure. By examination of serial sections at P14, cysts (asterisk) demonstrate a clear demarcation to the surrounding interstitial tissue. Numerous tubules (arrow) are located in close vicinity to the cysts, but no transition of a cyst to a tubule or any other renal structure is detectable.

**Supplemental Figure 2.**
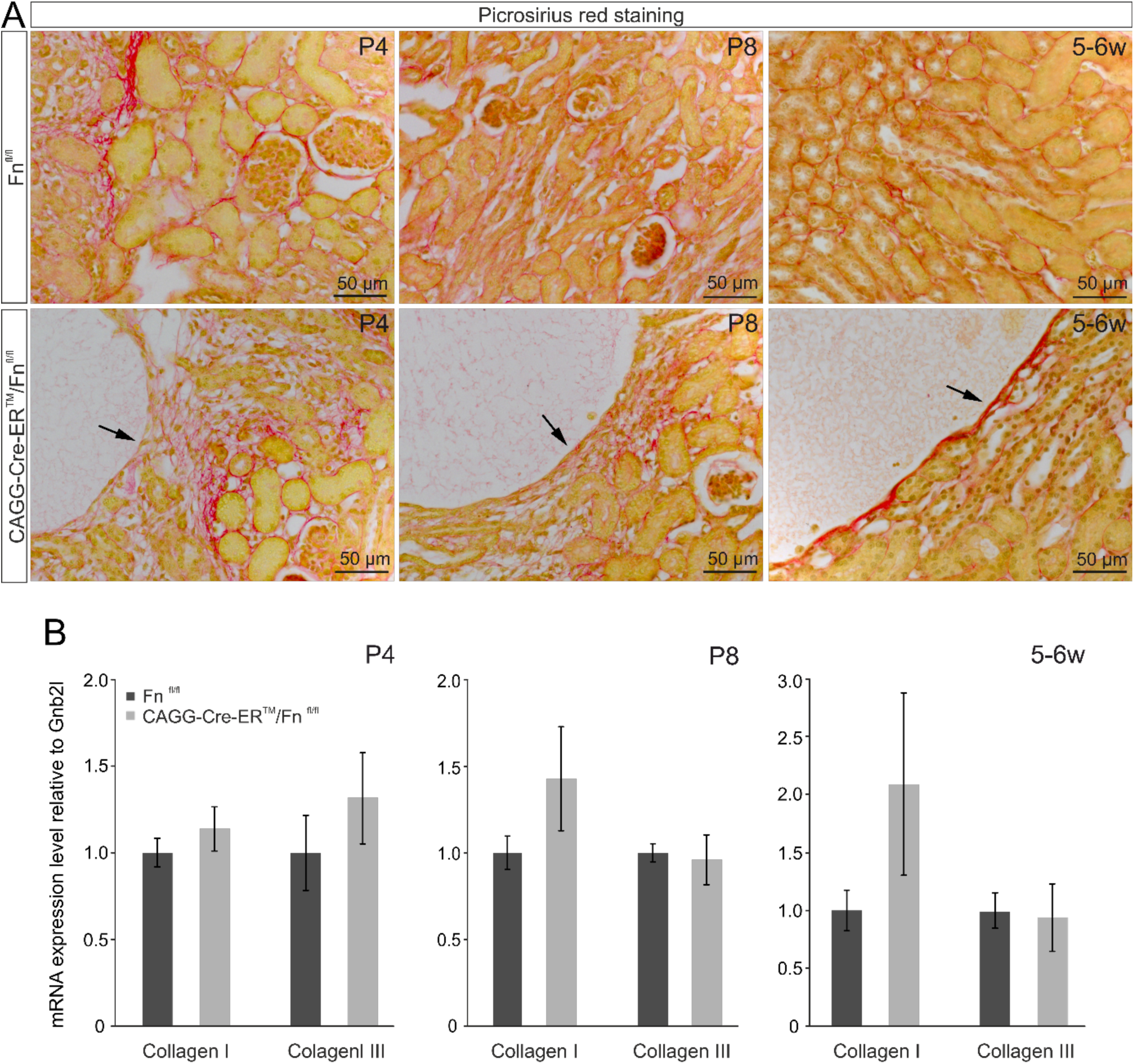
Increasing deposition of collagen fibers along the cysts. For induction, experimental and control mice were treated from P1 three times daily with tamoxifen eye drops (2.5 mg/ml). The experiments were conducted at P4, P8, and 5-6 weeks of age. **A**. Picro sirius red staining performed with frontal sections from P4 and P8 kidneys shows a characteristic distribution of collagen fibers throughout the interstitium of CAGG-Cre-ER^™^/Fn^fl/fl^ kidneys (left and middle lower panel), which is quite similar to that seen in the control Fn^fl/fl^ littermates (left and middle upper panel). At P4 and P8, Picro-Sirius red staining does not show an increased amount of collagen along the cysts (arrow). Whereas, at the age of 5-6 weeks there is a clear accumulation of collagen fibers along the cysts (arrow) in CAGG-Cre-ER^™^/Fn^fl/fl^ kidneys. **B**. Quantitative real-time RT-PCR analyses of whole kidney samples show a trend towards an increase in collagen I expression in the CAGG-Cre-ER^™^/Fn^fl/fl^ mice with increasing age, compared to the controls. In contrast, mRNA expression of collagen III does not appear to be affected by the deletion of Fn (n ≥ 4, n ≥ 7; mean ± SEM).

