## Supplemental Figure 1 for "Fibronectin deficiency in newborn mice leads to cyst formation in the kidney"

**Supplemental Figure 1. Tubule-independent cystogenesis in FN-deficient mice at P4 and P14.** Semi-thin serial sections of a renal cyst at P4 and P14. For induction, the P4 animals were treated three times daily with tamoxifen eye drops (2.5 mg/ml) on P1 and P2. P14 animals were treated three times daily with tamoxifen eye drops (2.5 mg/ml) starting from P1. Histological analysis of the kidney was performed at P4 or P14 by Richardson staining. Serial sections at P4 do not show a transition of the cyst (asterisk) to a renal structure. By examination of serial sections at P14, cysts (asterisk) demonstrate a clear demarcation to the surrounding interstitial tissue. Numerous tubules (arrow) are located in close vicinity to the cysts, but no transition of a cyst to a tubule or any other renal structure is detectable.

**Supplemental Figure 1**

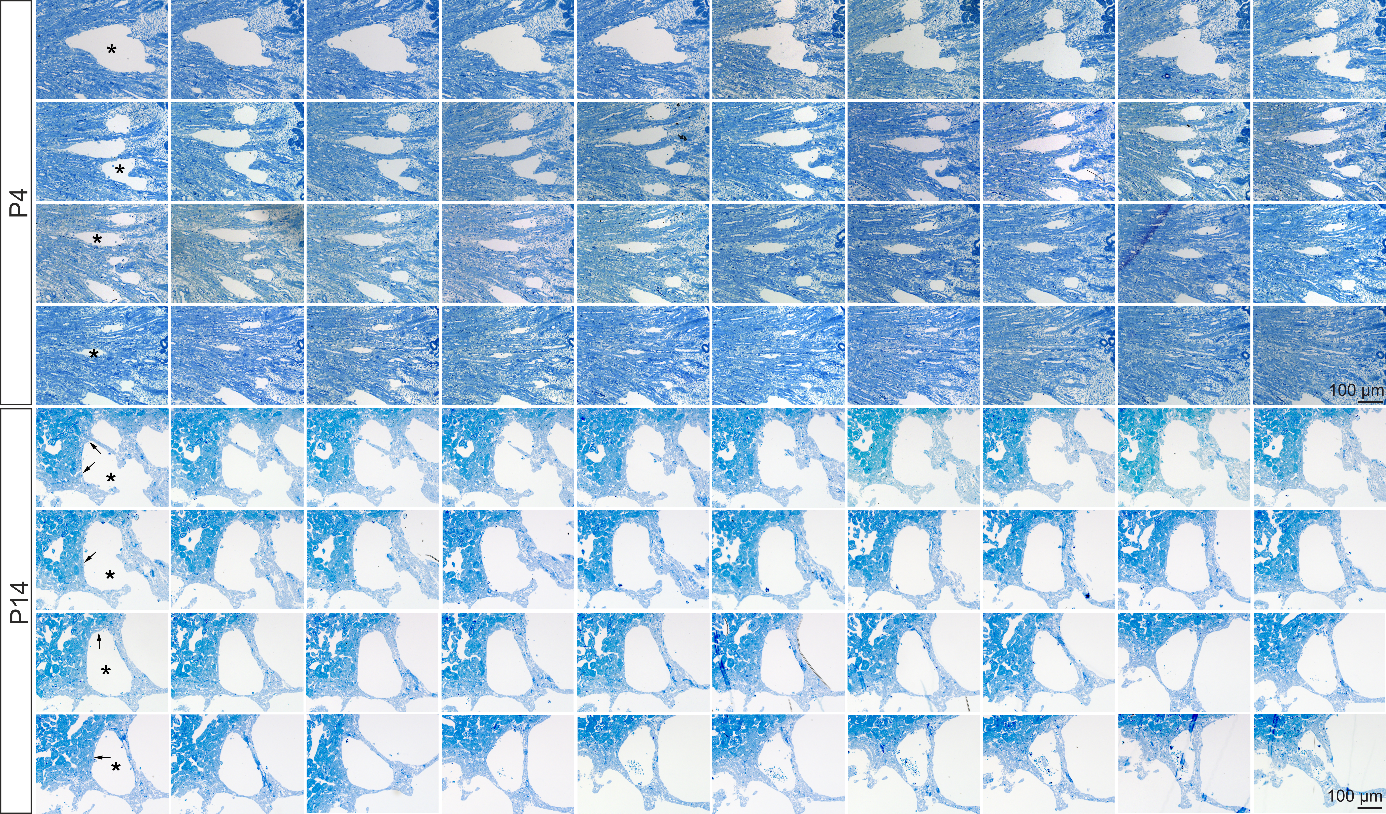
